# Emergence and organization of adult brain function throughout child development

**DOI:** 10.1101/2020.05.09.085860

**Authors:** Tristan S. Yates, Cameron T. Ellis, Nicholas B. Turk-Browne

## Abstract

Adult cognitive neuroscience has guided the study of human brain development by identifying regions associated with cognitive functions at maturity. The activity, connectivity, and structure of a region can be compared across ages to characterize the developmental trajectory of the corresponding function. However, observed developmental differences may not only reflect the maturation of the function but also its organization across the brain. That is, a function may be mature in children but supported by different brain regions and thus underestimated by focusing on adult regions. To test these possibilities, we investigated the presence, maturity, and localization of adult functions in children using probabilistic shared response modeling, a machine learning approach for functional alignment. After learning a lower-dimensional feature space from fMRI activity as adults watched a movie, we translated these shared features into the anatomical brain space of children 3–12 years old. To evaluate functional maturity, we correlated this reconstructed activity with the children’s actual fMRI activity as they watched the same movie. We found reliable correlations throughout cortex, even in the youngest children. The strength of the correlation in the precuneus, inferior frontal gyrus, and lateral occipital cortex increased over development and predicted chronological age. These age-related changes were driven by three types of developmental trajectories across distinct features of adult function: emergence from absence to presence, consistency in anatomical expression, and reorganization from one anatomical region to another. This data-driven approach to studying brain-wide function during naturalistic perception provides an abstract description of cognitive development throughout childhood.

**Significance Statement:** When watching a movie, your brain processes many types of information—plotlines, characters, locations, etc. A child watching this movie receives the same input, but some of their cognitive abilities (e.g., motion detection) are more developed than others (e.g., emotional reasoning). Beyond anatomical differences, when does the child brain begin to *function* like an adult brain? We used a data-driven approach to extract different aspects of brain activity from adults while they watched a movie during fMRI. We then predicted what the brain activity of a child would look like if they had processed the movie the same way. Comparing this prediction with actual brain activity from children allowed us to track the development of human brain function.

## Introduction

The advent of non-invasive neuroimaging techniques opened a new window into the study of human cognitive development. Initial studies of children examined anatomical brain regions associated with particular cognitive functions in adults, such as prefrontal cortex for executive function (Luna et al. 2001) and the amygdala for fear processing (Thomas et al. 2001). This approach was effective in characterizing the development of these brain regions. However, given that cognitive functions can be subserved by different brain regions at different ages (Schlaggar 2002; Brown et al. 2005; Nelson et al. 2003; Bayet and Nelson 2019; Durston et al. 2006; Jolles et al. 2011; Thomason et al. 2008), focusing on adult regions may underestimate the maturity of these functions in children. This is consistent with the *interactive specialization* framework (Johnson 2001, 2011), which emphasizes that brain regions specialize through network interactions and experience. One striking example is the development of visual object recognition: face processing is initially supported by both left and right fusiform gyrus, but as children learn to read, a region selective for visual words emerges in the left fusiform gyrus and face processing begins to shift to being right lateralized (Dundas et al. 2013; Centanni et al. 2018; Dehaene-Lambertz et al. 2018). In this case, development does not result from the maturation of brain regions in isolation, but rather from competition between regions.

The current study builds on this prior work in two ways. First, there are multiple interpretations for patterns of cognitive development in the brain such as those described above (Brown et al. 2006; Poldrack 2010). For example, a shift from distributed to focal processing over development (Durston et al. 2006) could be evidence of increased efficiency of brain regions, decreased reliance on other regions for “support,” changes in the computations being performed, or simply artifacts of greater variability in region localization in developing populations. We establish an unbiased, data-driven approach that can capture different kinds of developmental change under a common framework. Second, most brain-based studies of cognitive development pursue a standard lab approach of isolating cognitive functions. Although vital for manipulating and tracking exactly what is changing, an alternative, naturalistic approach could provide a more comprehensive and ecological sense of how the brain is developing. Indeed, in adult cognitive neuroscience, naturalistic paradigms such as movie-watching have yielded unexpected insights into how the brain processes information across time-scales and domains (Sonkusare et al. 2019). Given the challenge of collecting developmental data, such paradigms have the additional advantage of broadly sampling cognition without requiring multiple experiments.

To track the neural development of cognitive functions within and across brain regions, we applied functional alignment (Chen et al. 2015) to an open-access dataset of children aged 3–12 and adults watching a movie during fMRI (Richardson et al. 2018). We used open-source software for shared response modeling (SRM; Kumar et al. 2020) to extract temporal features of brain activity that were shared across the adults. These features can be thought of as capturing abstract cognitive functions that vary distinctively from each other across the movie in a way that is consistent in adults. We then mapped the children into this lower-dimensional feature space. These mappings were used in reverse to port adult fMRI activity into each child’s anatomical brain space. Comparing this reconstructed activity to the child’s actual fMRI activity allowed us to quantify the expression of adult cognitive functions throughout childhood. Although these abstract functions cannot be cleanly identified with specific psychological constructs, a key advantage of this data-driven approach is that they can be aligned across adults and children without making any anatomical assumptions. There is no requirement that functions are instantiated in the same brain regions across individuals, whether within or between ages. In fact, our approach would be equally sensitive to the development of functions that emerge within one region as to functions that reorganize from one region to another.

## Materials and Methods

### Data

The Partly Cloudy dataset was obtained from the OpenNeuro database (accession number ds000228). A full description of data acquisition can be found in the original paper (Richardson et al. 2018). Participants with neuroimaging data available consisted of 33 adults (18–39 years old; M = 24.8, SD = 5.3; 20 female) and 122 children (3.5–12 years old; M = 6.7, SD = 2.3; 64 female).

### fMRI acquisition and preprocessing

Participants watched an animated movie (Sohn and Reher 2009) that lasted approximately 5 minutes while undergoing fMRI. No explicit task was given beyond staying still and paying attention to the movie. Adults and older children used the standard Siemens 32-channel head coil. For younger children, one of two custom 32-channel phased-array head coils was used (smallest coil: *N*=3, *M*(*s.d.*)=3.91(0.42) years old; smaller coil: *N*=28, *M*(*s.d.*)=4.07(0.42) years old). fMRI data were collected using a gradient-echo EPI sequence (TR=2s, TE=30ms, flip angle=90 degrees, matrix=64×64, slices=32, interleaved slice acquisition) covering the whole brain. To correct for slight variations in the voxel size and slice gap parameters across participants, data were resampled to 3mm isotropic with 10% slice gap (the modal parameters). Children also participated in a number of behavioral tasks beyond the scope of the current study.

Preprocessing of the structural and functional MRI data was performed with fMRIPrep (v1.1.8; Esteban et al. 2019). First, T1-weighted structural images from an MPRAGE sequence (GRAPPA=3, slices=176, resolution=1mm isotropic, adult coil FOV=256mm, child coils FOV=192mm) were corrected for intensity non-uniformity using N4BiasFieldCorrection (v2.1.0) and skull-stripped using antsBrainExtraction.sh (v2.1.0, OASIS template). Surface reconstruction was performed by FreeSurfer (v6.0.1). Nonlinear registration for spatial normalization was performed with the antsRegistration tool (ANTs v2.1.0). Registrations were visually inspected and the quality of fit did not seem to differ across child and adult participants. Functional images were slice-time corrected using 3dTshift from AFNI (v16.2.07), then motion corrected using FSL’s mcflirt (v5.0.9). Co-registration to the structural scan was performed with 9 degrees of freedom using bbregister in FreeSurfer (v6.0.1). Transformations were concatenated using antsApplyTransform (v2.1.0). Physiological noise regressors (temporal and anatomical) were extracted using component-based noise correction and frame-wise displacement was estimated using Nipype.

### Experimental design

We used SRM (Chen et al. 2015; Turek et al. 2018) to identify activity in the developing brain that could be predicted from adult brain activity (illustrated in Figure 1). This method assumes that all participants were shown the same stimulus with the same number of time-points but does not require that they have the same number of voxels. First, the time-points from a group of adults were evenly split into training and test sets. We used one half of the adult data to learn the shared features space, consisting of features that captured shared temporal variance across adults, as well as the mappings between individual adults’ brain activity and this shared space. Prior to any other analyses, we ran this analysis on subsets of the adult data varying the number of features (5–80) and found that 10 features gave the highest whole-brain signal reconstructions across adults (M = 0.087, SD = 0.031).

**Figure 1:**
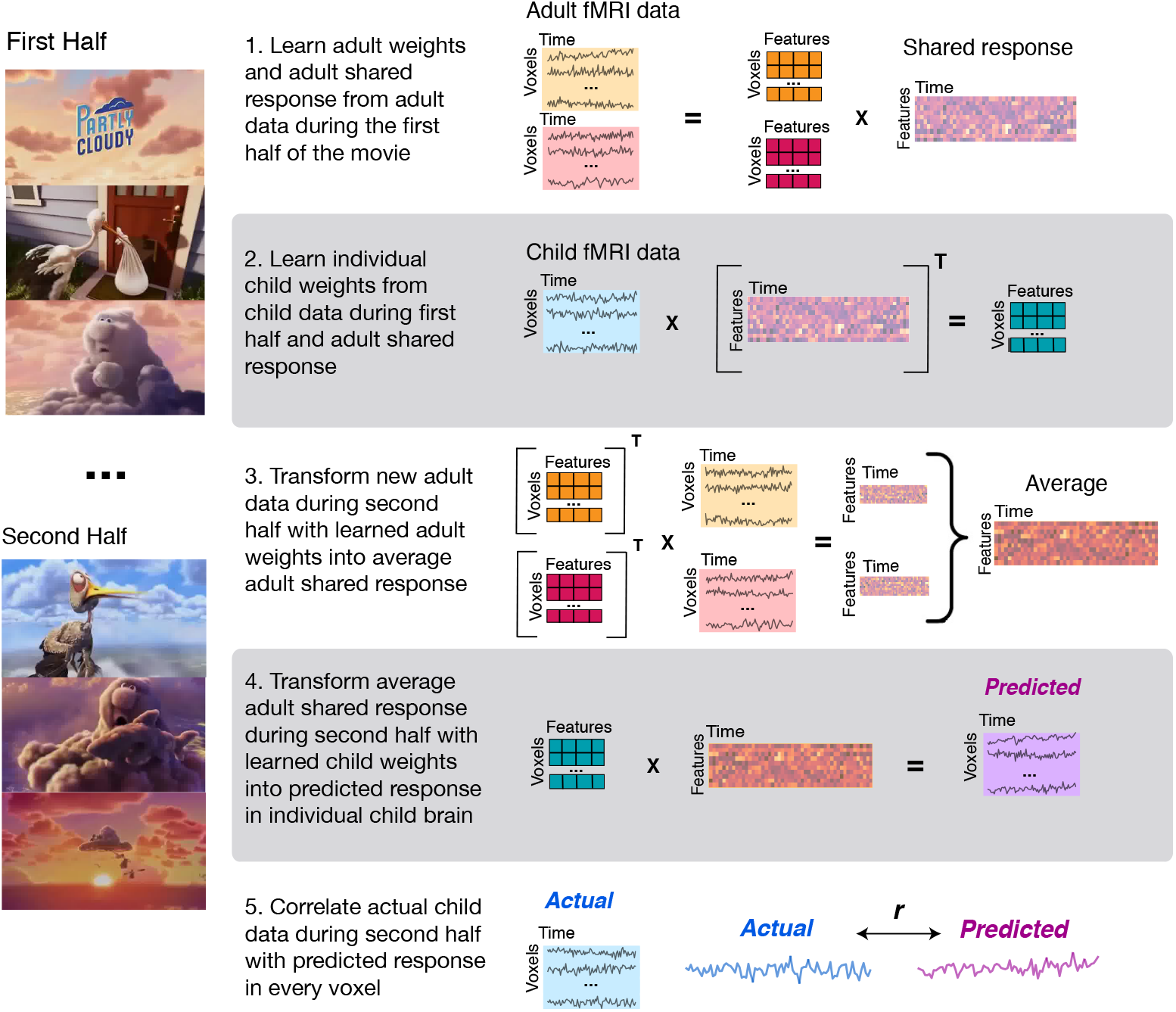
Schematic of the signal reconstruction pipeline. An SRM was trained on a group of adults (N=33) and then each of the children (N=122) was fit into this space. Adult data not used to train the model were transformed into the shared space and averaged. This was then projected into each child’s brain and correlated with their actual activity.

After learning 10 shared features in adults using one half of the adult data, we found the mapping (voxels by features) between an individual child’s functional activity (voxels by time) and the shared response (time by features) for this same portion of the movie. Singular value decomposition was implemented to solve for the orthogonal weight matrix. Values in each cell of this resulting weight matrix denote how strongly a given voxel in the child expresses each of the 10 features discovered from the adult data. Next, we used the remaining half of the adult data to quantify how the 10 shared features were expressed in data not involved in SRM training. Each adult’s transposed weight matrix (features by voxels) was used to transform their raw voxel activity (voxels by time) into the shared feature space (features by time). We then averaged these shared responses across all adults to find the canonical adult response in terms of shared features during this part of the movie. Finally, each child’s weight matrix (voxels by features) was used to transform the average adult shared response (features by time) into the child’s brain space (voxels by time). This predicted response represents what the child’s brain activity would look like if they expressed the same shared features of adults. We quantified the extent to which this was true by correlating the child’s actual raw response with this predicted response for each voxel separately. Thus, higher signal reconstruction reflected greater adult-child functional similarity — i.e., more adult-like functions in the child’s brain — agnostic to the anatomical localization of these functions in either group. The voxelwise map of predicted-actual correlations for each child was averaged across individuals within age groups. We ran this entire procedure twice, training the SRM on the first half of the movie and testing on the second half in one fold, and then vice versa in another fold, and present results averaged across these folds.

Our first objective was to assess the degree to which children’s brain activity could be reconstructed from shared features learned in adults. We quantified the noise ceiling for this group-level signal reconstruction by leaving one adult participant out of SRM training, correlating that individual’s predicted and actual brain activity, and then iterating through each adult. This was treated as the noise ceiling because the held-out participant was from the same age group used to train the SRM. Our second objective was to quantify how signal reconstruction may change over development, and whether this could be a useful measure for predicting an entirely held-out child’s age. Finally, we explored how the individual features that comprise the shared response may exhibit different developmental trajectories throughout childhood.

### Statistical analysis

We used bootstrap resampling methods to statistically evaluate our results non-parametrically (Efron and Tibshirani 1986; Kim et al. 2014; Fan et al. 2020). For each effect of interest, at the last step of the analysis we randomly sampled participants with replacement 10,000 times, averaging the effect across the sample of each iteration. The logic of this approach is that if an effect is reliable across participants, the participants should be interchangeable, and a similar group effect should be observed in each iteration. The resampled values across all iterations reflect a sampling distribution of the effect of interest, further providing confidence intervals on the original effect. Null hypothesis testing can be performed by determining the proportion of resampled values that were of the opposite sign as the original effect. The original effect can also be normalized into a *z* statistic by dividing the mean of the resampled distribution by its standard deviation. For voxelwise analyses, this was performed in each voxel to create a statistical map. This map was corrected for multiple comparisons using a cluster-based correction in FSL’s cluster tool (cluster-forming threshold, *p<*0.001). Corrected p-values were found using Gaussian Random Field Theory and the smoothness estimated from the original map.

We quantified the relationship between signal reconstruction and age by first fitting a linear regression model for each voxel. We then used the same bootstrap resampling approach described above, now resampling participants who contributed to the relationship in each iteration. We calculated the *p*-value as the proportion of iterations on which the correlation coefficient from the linear regression model went in the opposite direction from the original model. We compared this model against other types of models that have been used previously in developmental cognitive neuroscience (Schlichting et al. 2017). For each voxel, we fit five regression models: (1) a linear model with age alone as the predictor (as above), (2) a linear model with age and sex as predictors, (3) a linear model with age and sex as predictors plus an age-by-sex interaction term, (4) a quadratic model with just age as a predictor, and (5) a quadratic model with age and sex as predictors plus an age-by-sex interaction term. We found that the linear model with age fit best (Table 1).

**Table 1:** Different models used to predict signal reconstruction. For each voxel, the model with the lowest AIC was assigned to that voxel. The average and standard deviation of AIC values for a given model for voxels where that model was the best is shown in the right column. Overall, the linear model used in our main analyses with age as the only predictor best described the data.

| Regression Model | Number of Voxels | Average AIC |
| --- | --- | --- |
| $SigRecon \sim Age$ | 39111 | 2.41 (0.56) |
| $SigRecon \sim Age + Sex$ | 60 | 5.21 (1.39) |
| $SigRecon \sim Age + Sex + Age * Sex$ | 31 | 4.76 (1.91) |
| $SigRecon \sim Age^2$ | 49 | 3.75 (2.01) |
| $SigRecon \sim Age^2 + Sex^2 + Age * Sex$ | 31 | 3.71 (1.05) |

We used leave-one-out cross-validation to predict the age of children from signal reconstruction. For each iteration, we fit the linear regression model between signal reconstruction and age in a training set of *N*-1 participants. We did this separately for each voxel and retained the clusters that were significant within the training set (based on the previously described bootstrap resampling method). Note that we ignored the sign of the significant relationship in a cluster and thus it was possible to find negative beta values. We then fit a regularized ridge regression (penalty = 1) across voxels from the significant clusters. To predict age in the held-out *N ^th^* test participant, we input their signal reconstruction scores across these voxels and output an estimated age. Finally, we calculated the Pearson correlation and mean-squared error between the chronological and predicted ages of children across iterations.

In addition to reconstructing all 10 adult features in children, we also performed signal reconstruction for individual adult features. To test single-feature reconstruction within the adults, we could not perform the fully cross-validated approach described above of leaving one adult participant entirely out of both the training set used to learn the SRM and the testing set used to generate predicted activity. This is because each training set would have contained a unique set of adults, which could lead to different features and/or a different ordering of features in the shared space. We would therefore not be confident that we were considering the same feature across folds. Instead, we included all adults when training the SRM on one half of the movie, so that there would be a consistent shared space across adults and as used to reconstruct children. Nevertheless, we left one adult out when averaging the adult shared response for the other half of the movie, using the expression of the selected feature in all but that adult to predict their neural activity. Because the reconstructed adult was used in SRM training, we included a 10 time-point buffer between their training and test data to minimize non-independence. Signal reconstruction of individual features in children was identical to the main analysis, except based on separate weight matrices for each child mapping from their voxel space to a given adult feature. That is, although the adult data could not be fully cross-validated, the data from children remained completely untouched during SRM training.

In the single-feature analysis, we sought to quantify how brain regions changed in their expression of features over development. We thus defined regions of interest using the Schaefer brain atlas (Schaefer et al. 2018). This atlas consists of 100 parcels discovered from resting-state connectivity data in adults and matched to 17 functional networks (Yeo et al. 2011). We reconstructed each of the ten features in adults and children and then calculated the average signal reconstruction scores across voxels in each of the 100 parcels. For statistical analysis, we used the same bootstrap resampling procedure across the participants in a given age group, separately for each parcel and feature. To correct for multiple comparisons across the parcels, we used Bonferroni correction (100,000 bootstrap iterations were run to gain precision on *p*-values for thresholding). Finally, the regions that survived correction were ranked according to the strength of signal reconstruction.

### Code accessibility

The analysis code for running the signal reconstruction analysis pipeline will be made available on Github.

## Results

### Adult-like brain function in early to middle childhood

We first characterized how well adult brain activity could be reconstructed from other adults. Signal reconstruction in adults was widespread throughout the brain, especially in occipital and parietal cortices (Figure 2A). This indicates that the shared features learned by SRM can account for a good proportion of adult brain activity during a short movie. Remarkably, signal reconstruction was also widespread in the children despite the fact that the functional features were defined entirely in adults (Figure 2B). In the child brain, adult functions were most strongly represented in lateral occipital and posterior medial regions, albeit weaker than in the adult brain.

**Figure 2:**
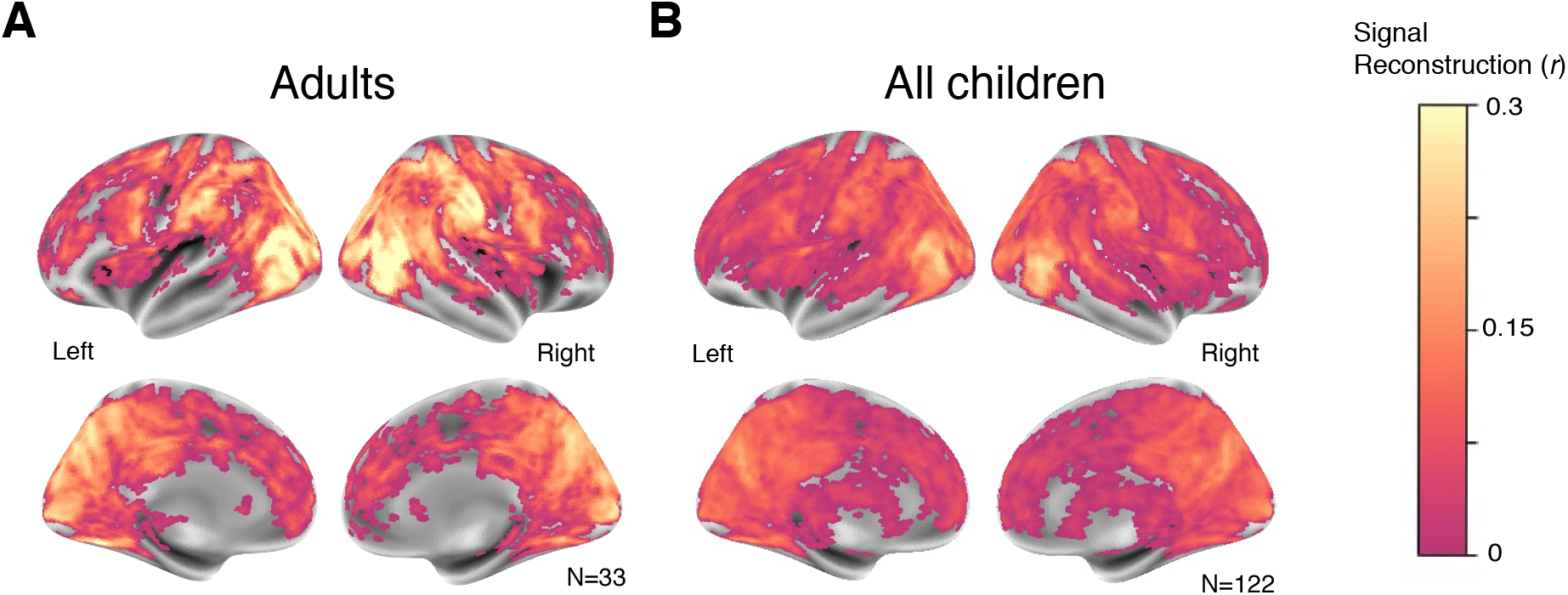
Reconstruction of adult function in children. In all brain plots, the strength of signal reconstruction is denoted by color and only regions that survived cluster correction are plotted. (A) Signal reconstruction for a group of adults predicting an independent adult’s functional activity is reliable throughout much of the brain. (B) Signal reconstruction remains reliable (though weaker) for adults predicting functional activity in children ranging from 3–12 years old.

### Relationships between age and signal reconstruction

The previous analysis collapsed across all children, but the extent and location of signal reconstruction may vary with age. We quantified these relationships by correlating signal reconstruction in each voxel with chronological age across children. After correcting for multiple comparisons, adult function was positively correlated with age in regions including the bilateral precuneus, bilateral lateral occipital cortex, postcentral gyrus, and inferior frontal gyrus (Figure 3a). No regions showed a reliable negative correlation. Alternative models taking into account children’s sex and testing for quadratic relationships did not generally provide better fits than this linear model (Table 1). The basic linear model with age alone gave the lowest Akaike information criterion (AIC) values for the majority of voxels, and therefore minimized the information loss when trading off with model complexity. Furthermore, individual voxels in which other models had the lowest AIC values were scattered across the brain, suggesting that they were capturing noise and providing further evidence that the basic linear model performed best.

**Figure 3:**
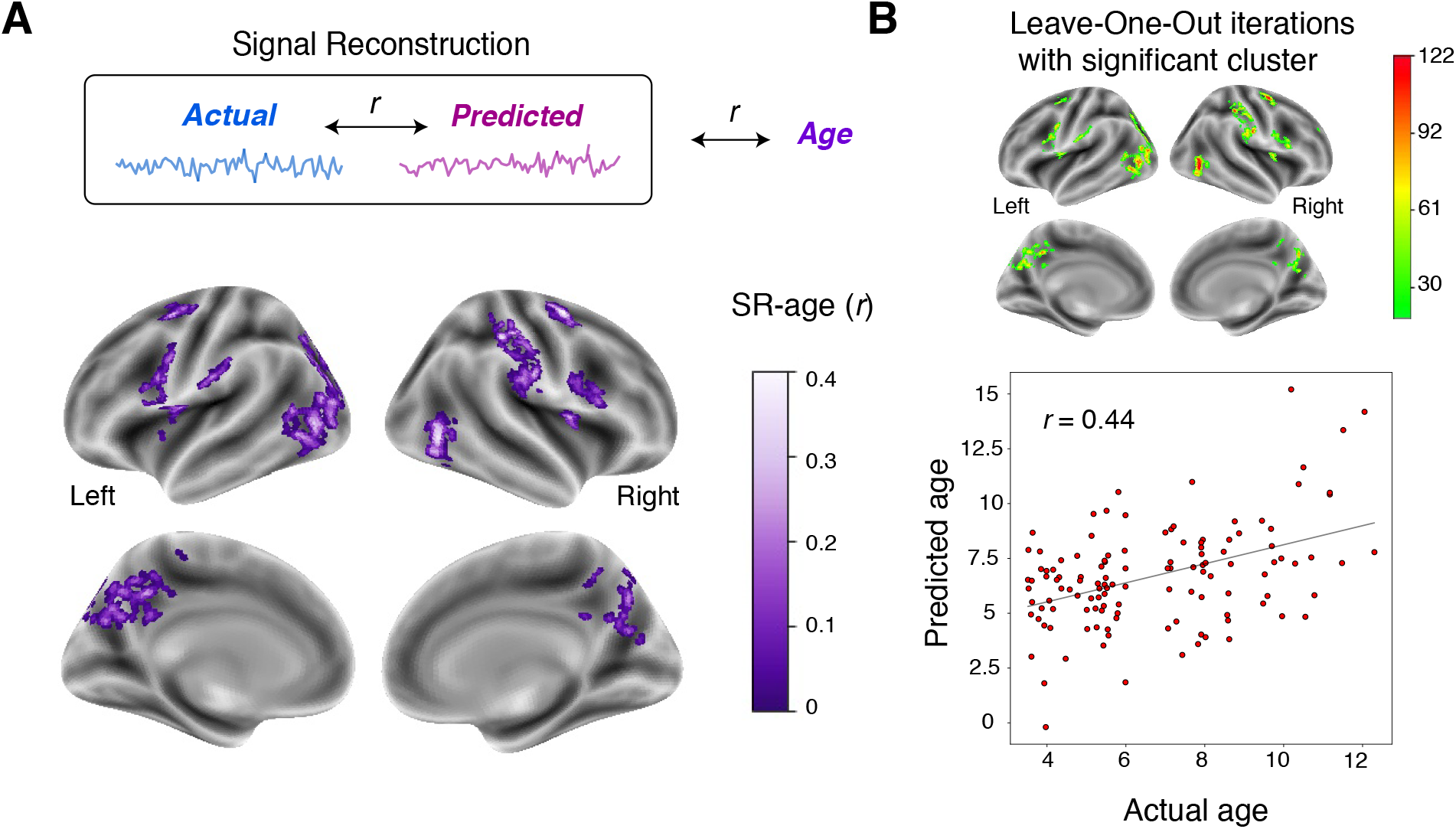
Relationship between signal reconstruction and age. (A) Brain regions with a reliable correlation between signal reconstruction and age across all 122 children, colored by signal reconstruction. (B) Similar regions are found in leave-one-child-out iterations of the age prediction analysis. Colors closer to red signify regions that were significant in most of the iterations. Using signal reconstruction scores from these regions, we were able to accurately predict the held-out child’s chronological age.

### Out-of-sample prediction of a child’s age from signal reconstruction

With chronological age related to signal reconstruction in several regions of the brain, it may also be possible to predict the age of a previously unseen child. In a nested cross-validation analysis, we first trained a linear regression model between signal reconstruction and age for each voxel in all but one child. Blind to this child, we determined which voxels showed a significant relationship with age again through bootstrapping and cluster correction. We then trained a ridge regression model on these significant voxels. This model was used to predict the held-out child’s chronological age from their multivariate pattern of signal reconstruction scores across the voxels. This procedure was repeated 122 times to use each child as the held-out test data once. Note that the significant clusters varied slightly across iterations because the training set changed when different children were used as test data. Finally, we correlated the predicted and actual ages (Figure 3B), and found a strong relationship (*r*=0.436, *p<*0.001). Indeed, our model had a mean-squared error of 6.05, meaning that our average error in age prediction was 2.46 years across an age range of 8.78 years.

### Reliable signal reconstruction in all age groups

The relationship between signal reconstruction and age could reflect a lack of adult function in early childhood that emerges in middle childhood. To evaluate this possibility, we divided children into five age bins (3.5–4.5, 4.5–5.5, 5.5–7.5, 7.5–9.5, 9.5–12.3 years old), each containing roughly the same number of participants (*N*=20–26). This was done for analytical convenience and was not intended to suggest discrete developmental stages. Although signal reconstruction increased with age, we nevertheless found reliable signal reconstruction in every age group. This includes lateral occipital, posterior medial, and supramarginal regions, even in the youngest children aged 3–5 (Figure 4). Signal reconstruction emerged in frontal regions around age 5, and became more pronounced in the older groups. To obtain a global measure of adult-child similarity, we correlated the unthresholded maps of signal reconstruction for each age group with that of adults. There was reasonable agreement in all groups, though the amount of variance explained grew from 25% in the youngest children to 64% in the oldest children.

**Figure 4:**
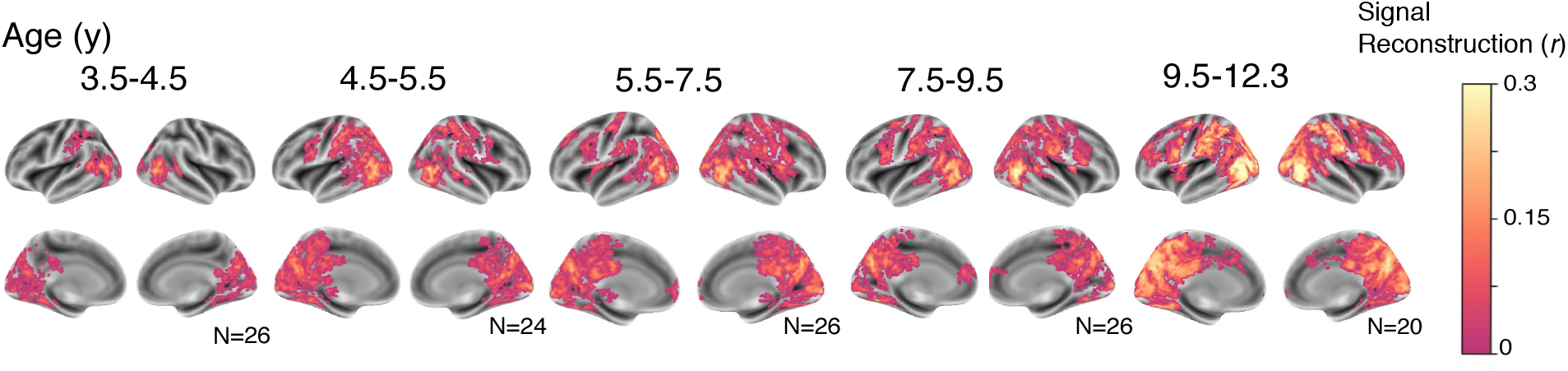
Signal reconstruction of adult features was reliable even in the youngest children, but spread anatomically and grew in strength throughout childhood. To quantify this developmental change, we correlated the unthresholded voxelwise signal reconstruction in each age group with that of adults, revealing increasing maturity: 3.5–4.5 years, *r*=0.507; 4.5–5.5 years, *r*=0.587; 5.5–7.5 years, *r*=0.601; 7.5–9.5 years, *r*=0.646; 9.5–12.3, *r*=0.797).

### Controlling for age-related noise in signal reconstruction

Increases in signal reconstruction over development may result from younger children being “noisier” than older children and adults, including because of differences in task compliance, preprocessing quality, and/or BOLD physiology (Phan et al. 2018; Harris et al. 2011). Children did move their heads more than adults overall, but this did not track with age across children (analysis of number of time-points exceeding 2-mm motion threshold from Richardson et al. 2018): one-way ANOVA across age groups, *F*(4,116)=1.162, *p*=0.331; correlation with age across children, *r*=-0.112, *p*=0.219.

Moreover, we can estimate and control for noise in different age bands using the noise-ceiling approach from adults (Figure 2A). For each age group, we held one child out and used SRM to learn shared features in the remaining children of that group. We then predicted the held-out child’s voxel activity, correlated it with their actual activity, and averaged across significant clusters to derive a global within-group signal reconstruction score for each child. The average score across children in a group provides a measure of the reliability of functional brain activity in that group. This within-group signal reconstruction correlated with chronological age (*r*=0.359, *p<*0.001), consistent with decreasing noise over development, and with adult-group signal reconstruction (*r*=0.647, *p<*0.001). Critically, however, the correlation between adult-group signal reconstruction and chronological age (*r*=0.417, *p<*0.001) persisted after controlling for within-group signal reconstruction (*r*=0.261, *p*=0.003). In contrast, the correlation between within-group signal reconstruction and chronological age did not hold after accounting for adult-group signal reconstruction (*r*=0.128, *p*=0.158). These results suggest that adult features capture more about child brain function than changes in noise over development.

### Emergence and reorganization of adult function over child development

There are at least two other potential explanations for the age-related increases in signal reconstruction we observed. First, the adult function being reconstructed may be completely absent from younger children and mature over development to become present in older children (emergence). Second, the adult function may be present in both younger and older children but expressed in different brain regions over development (reorganization). By both accounts, the brain regions expressing the function in older children would not express it as strongly in younger children. The accounts differ, however, in that reorganization but not emergence predicts that the function would be expressed in other brain regions in younger children.

Emergence and reorganization are difficult to distinguish with the analysis approach used so far. By predicting the activity of each voxel as a weighted combination of all adult features, we may have obscured developmental trajectories that differed across features. We thus modified our pipeline to predict activity in children from individual adult features, each of which captures a narrower, more unique range of adult function. The modification occurred in step 4 (Figure 1), where we now transformed only one average adult feature at a time into the voxel space of a child. Individual features do not necessarily isolate single functions, and emergence and reorganization are not mutually exclusive, so it may be possible to observe both patterns within a feature. We used a functional atlas (Schaefer et al. 2018) to identify regions that showed the strongest signal reconstruction for a given feature per age group.

We found evidence of both emergence and reorganization across different adult features, as well as a third pattern in which a feature was expressed in the same brain region(s) across development (consistency). Representative features illustrating these three types of trajectories are depicted in Figure 5 (for all features, see Figure 5-1). For example, Feature 4 was not reliably expressed in the two youngest age groups and emerged in the lingual gyrus of older children and adults (Figure 5A). In contrast, Feature 6 was expressed most strongly in the posterior cingulate and lingual gyrus consistently throughout development (Figure 5B). Finally, Feature 7 was expressed most strongly in the precuneus and posterior cingulate of children and migrated to different parietal and occipital regions in adults (Figure 5C). By measuring functional profiles regardless of anatomy, signal reconstruction revealed developmental changes both within and across brain regions.

**Figure 5:**
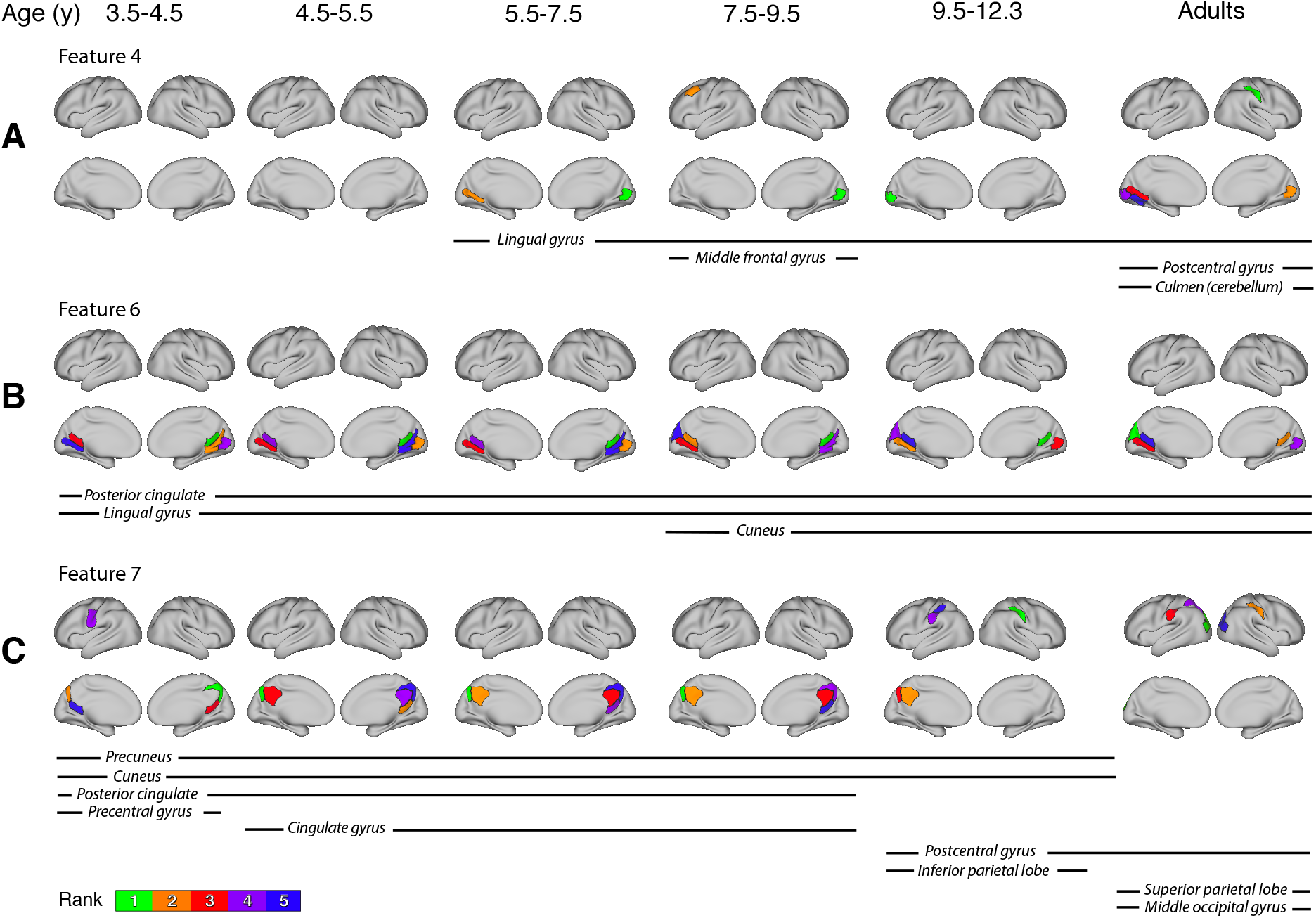
Trajectories of functional development within and across brain regions. (A–C) To understand the nature of developmental changes in signal reconstruction, we predicted activity from one adult feature at a time rather than all features. We used a functional parcellation to identify which regions expressed a given feature most strongly in each age group. Regions with significant signal reconstruction (*p<*0.05, corrected) were ranked by the strength of the reconstruction. For ease of visualization, here we color up to the top five regions for each feature and age group. The anatomical labels for these regions were obtained from the Talairach atlas. Three example adult features are depicted across ages, illustrating developmental trajectories we refer to as emergence (Feature 4), consistency (Feature 6), and reorganization (Feature 7). The remaining features are depicted in Figure 5-1.

## Discussion

In this study, we sought to bring a new perspective to the long-standing question of how and when the developing brain becomes “adult-like” (Somerville 2016; Johnson 2011). The typical approach for answering this question is to align children and adults into a common anatomical space and compare activity between groups in the same brain regions (Gogtay et al. 2004; Fair et al. 2009; Dosenbach et al. 2010; Cantlon and Li 2013; Richardson et al. 2018). Thus, even when the goal is to understand functional similarities and differences over development, anatomy serves as a guide and constraint. The alternative approach we employed is to align children and adults into a common *functional* space, which allowed us to quantify adult-like brain activity in children without making any assumptions about a consistent mapping between function and anatomy over development. In children as young as 3.5 years old watching a short movie, we found regions of the brain, especially in occipital cortex, that reliably expressed functional features shared amongst adults who watched the same movie. Based on where and how strongly these features were expressed, we were able to build a predictive model of age that depended only on brain activity during movie watching. Finally, we demonstrated the power of functional alignment by revealing features of adult function that emerge and reorganize across anatomical locations over development.

We interpreted increasing signal reconstruction with age as evidence of functional specialization and maturation in the developing brain. A related but slightly different framing is that brain functionality itself was not always changing in these cases, but rather it was the way that children deployed this functionality during the movie. For instance, if older children attended to the content of the movie in a more adult-like fashion than younger children, this may have affected perceptual input to downstream functions and increased similarity to adult brain activity. The defining characteristic of this interpretation is that younger children may possess the *capacity* for such functions but not engage them because of attentional differences in perceptual input. Even if attention was allocated similarly across age, richer schematic knowledge in older children may have enhanced their understanding of the movie narrative (Ghosh and Gilboa 2014; Brod et al. 2017) by highlighting connections between objects and events that may not otherwise be easily integrated. Again, younger children may have the capacity for this kind of integration in principle but be unable to deploy it without access to the relevant conceptual knowledge. Of course, attention and memory are functions of the brain, and so developmental differences in these functions are what we sought to characterize in the first place. The key point is that increasing signal reconstruction could reflect the development of a function or of necessary perceptual or conceptual precursors to that function.

A variant of the signal reconstruction approach allowed us to identify different types of developmental trajectories across adult features. Yet, these data-driven features remain abstract and are not easily decomposed into specific cognitive functions. Moreover, although the features captured unique and substantial variance shared across adults, each may still embed multiple cognitive functions with similar temporal profiles of brain activity. This has implications for interpreting features showing anatomical reorganization of function over development (e.g., Figure 5C). Specifically, our definition of reorganization was that the same functions were subserved by different brain regions over development — that is, a cognitive function that manifests in region X of younger children is expressed in region Y of adults. This could occur if the original region was co-opted by a different function (Behrmann and Plaut 2015) or if the nature of the function changed with increasing skill and expertise (Johnson 2001).

However, the possibility of multiple functions being embedded in a given feature suggests an alternative interpretation. Namely, these functions may have stable organization over development, but the relative weighting of the functions as captured by the feature may change. Consider a hypothetical feature that is active during the title and credits of the movie. This feature might capture multiple language-related functions engaged by these scenes, such as letter recognition in region X and semantic comprehension in region Y. We would expect even the younger age groups to respond to the orthography of the words and thus show signal reconstruction in X, but perhaps only the older children and adults would respond to the meaning of the words and show signal reconstruction in Y. Disentangling these possibilities requires a better understanding of how the abstract features from SRM relate to the contents of a movie and to the cognitive functions that are engaged. Future studies could make progress in this direction by selecting or designing movies to target specific cognitive functions. Indeed, a limitation of our study, and SRM more generally, is that the features extracted depend on the movie. For example, the short, silent cartoon we used likely engaged functions related to visual processing, object recognition, and narrative comprehension, and thus the signal reconstruction and developmental trajectories we observed would be limited to those functions. The use of other data for functional alignment, including from more naturalistic videos, different sensory modalities, or synchronized trials of varied cognitive tasks would sample cognition more broadly. This might allow SRM to learn a richer functional space that provides a more complete picture of functional brain development across childhood.

We focused on brain development, but the techniques in our paper could be applied productively to a number of questions that involve comparing functional activity across groups. For instance, learning the functional features shared amongst a clinical population and then reconstructing these features in an undiagnosed individual may be useful for predicting whether the individual will develop the condition. This method could also be used to assess how and when a learner’s brain starts to resemble that of an expert over the course of training. Because signal reconstruction does not require that the group and individual have the same brain sizes or even anatomical organization, this approach could even be applied between humans and non-human animals to trace how cognitive functions are shared over phylogeny. Indeed, there is no requirement that the group and individual be brains at all, which could, for example, allow states of a computational model to be ported into the brain for model-based analysis, or vice versa for brain-computer interfaces.

## Acknowledgements

T.S.Y was supported by NSF Graduate Research Fellowship DGE 1752134. N.B.T-B. was supported by NIH R01 MH069456, NSF CCF 1839308, and the Canadian Institute for Advanced Research. We are grateful for the effort and generosity of Hilary Richardson and the rest of the research team that originally collected this dataset in Rebecca Saxe’s lab at MIT. We also thank lab members for helpful feedback on earlier versions of the project.

## Extended Data

**Figure 5-1:**
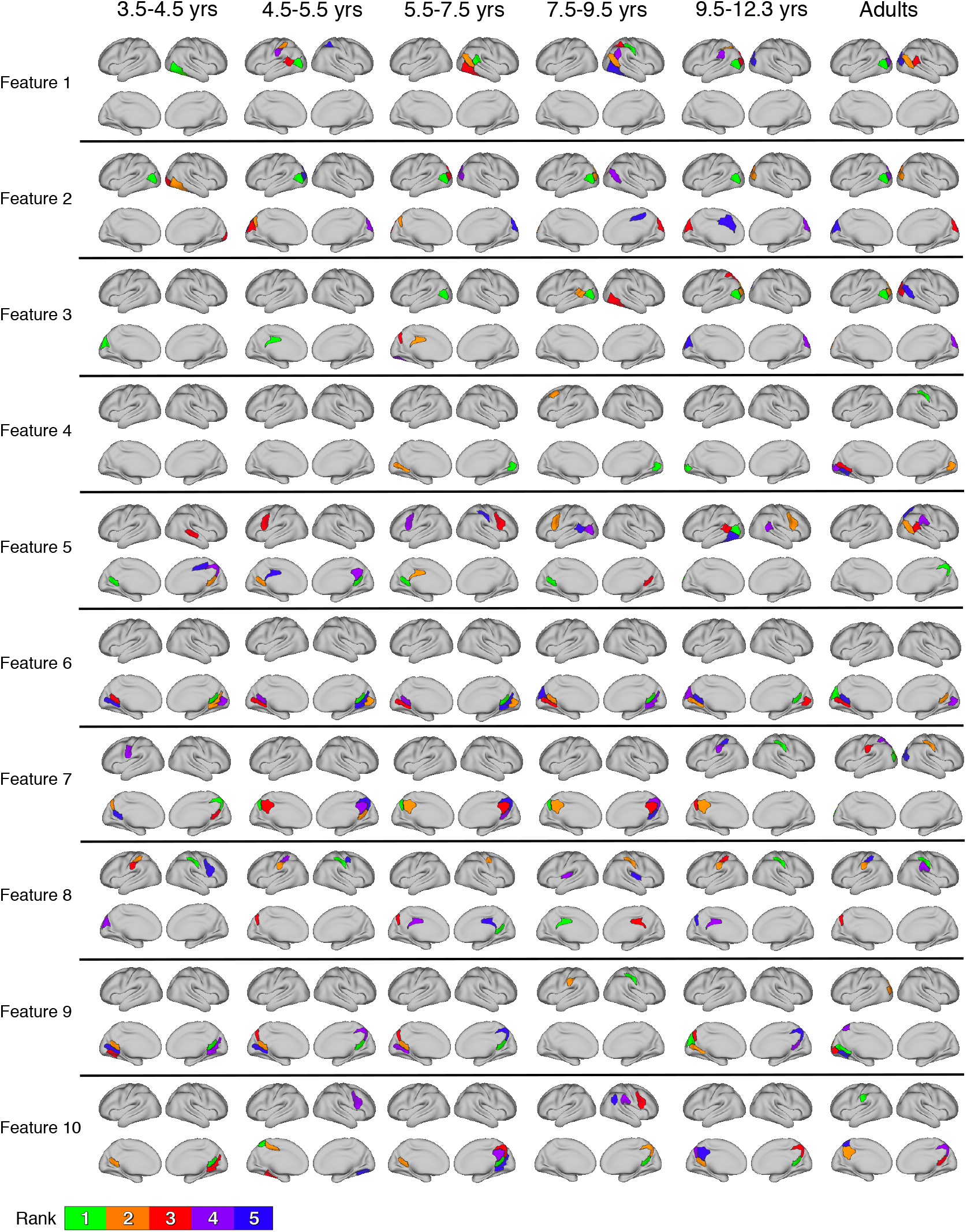
The top-5 significant parcels (*p<*0.05, corrected) from separate signal reconstruction of each adult feature in every age band.

